# Perineuronal nets and the neuronal extracellular matrix can be imaged by genetically encoded labeling of HAPLN1 *in vitro* and *in vivo*

**DOI:** 10.1101/2023.11.29.569151

**Authors:** Sakina P. Lemieux, Varda Lev-Ram, Roger Y. Tsien, Mark H. Ellisman

**Author notes:** Deceased August 24th, 2016 (Roger Y. Tsien). Correspondence: Sakina P. Lemieux, PhD Department of Neurosciences University of California, San Diego, Center for Research in Biological Systems 9500 Gilman Drive, MC 0608, La Jolla, CA 92093-0608. The authors report no conflict of interest.

## Abstract

Neuronal extracellular matrix (ECM) and a specific form of ECM called the perineuronal net (PNN) are important structures for central nervous system (CNS) integrity and synaptic plasticity. PNNs are distinctive, dense extracellular structures that surround parvalbumin (PV)-positive inhibitory interneurons with openings at mature synapses. Enzyme-mediated PNN disruption can erase established memories and re-open critical periods in animals, suggesting that PNNs are important for memory stabilization and conservation. Here, we characterized the structure and distribution of several ECM/PNN molecules around neurons in culture, brain slice, and whole mouse brain. While specific lectins are well-established as PNN markers and label a distinct, fenestrated structure around PV neurons, we show that other CNS neurons possess similar extracellular structures assembled around hyaluronic acid, suggesting a PNN-like structure of different composition that is more widespread. We additionally report that genetically encoded labeling of hyaluronan and proteoglycan link protein 1 (HAPLN1) reveals a PNN-like structure around many neurons *in vitro* and *in vivo*. Our findings add to our understanding of neuronal extracellular structures and describe a new mouse model for monitoring live ECM dynamics.

## Introduction

Neuronal long-term synaptic plasticity requires both intracellular and extracellular changes that contribute to memory persistence in the brain over time ^1–4^. ECM is present throughout the CNS; it protects neurons and synapses, maintains tissue integrity, and contributes to homeostasis and communication in ways that are not yet completely understood ^5^. ECM structures can be durable and contain molecules that are especially long-lived, persisting through normal protein turnover and CNS changes that occur during development ^6,7^. The ECM is composed of assorted proteoglycans including chondroitin sulfate proteoglycans (CSPGs), HAPLNs, and Tenascin R, all of which are produced and secreted in the developing CNS and remain relatively unorganized until critical period closure when they crosslink to form a net-like structure around cells ^8^.

ECM functions both in general structural support in the CNS and more directly in synapse formation, signaling, and stability. It controls plasticity by restricting excessive synaptogenesis and remodeling in adult mammals ^9^. Chemically, negative charges on CSPGs influence proximal ion and growth factor concentrations in the extracellular space, and localized matrix metalloproteinase (MMP) secretion during synapse strengthening can erode these structures by cleaving CSPGs at defined sites within the core protein, creating space for synaptic molecules to accumulate and exposing cryptic motifs that influence cell signaling ^10^. Studies using the bacterial-derived enzyme chondroitinase ABC (ChABC) to digest specific polysaccharides attached to proteoglycans *in vivo* have erased long-term memories, reopened developmental critical periods, and enhanced wound repair and synaptic plasticity in animal models ^11,12^.

PNNs are a specific form of neuronal ECM that surrounds the soma and proximal dendrites of parvalbumin (PV)-positive inhibitory interneurons in the CNS, and are important in memory encoding and retention ^9, 13^. Similar to broader ECM, traditionally defined PNNs are composed of CSPGs, HAPLNs, and Tenascin R, organized around hyaluronic acid (HA) chains that are anchored to the cell surface by the transmembrane hyaluronan synthase (HAS) (Fig 1A) ^3,14,15^. Camillo Golgi and Ramon y Cajal first defined this pericellular structure around certain neurons, and speculated that they may insulate neurons from the surrounding environment ^16^.

**Figure 1.**
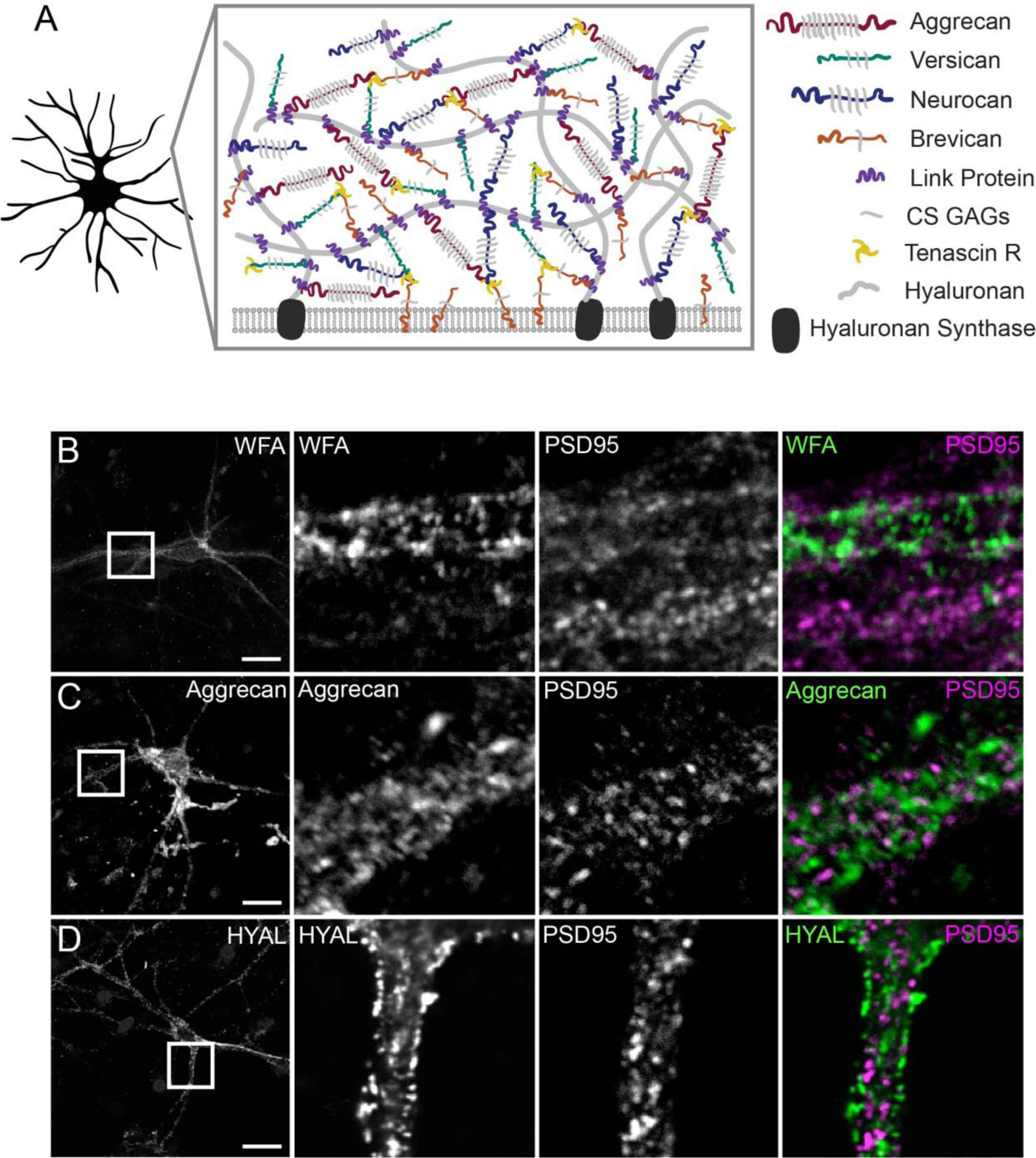
PNN labeling methods reveal an extracellular structure excluded from mature synapses in cultured neurons. ***A***, Perineuronal net structure and molecular composition. ***B***, Labeling with fluorescein-conjugated WFA and anti-PSD95 antibody. ***C***, Labeling with anti-aggrecan and anti-PSD95 antibodies. ***D***, Labeling with biotinylated HABP and anti-PSD95 antibody. Scale bars in first column, 20 μm; scale bars in second column, 5 μm.

PNN imaging in recent years has shown distinctive fenestrated structures that wrap around neurons and have holes filled by mature synapses ^17–19^. PNN has been widely imaged using plant-derived *Wisteria floribunda* agglutinin (WFA) and *Vicia villosa* agglutinin (VVA); these lectins tightly bind to *n*-acetylgalactosamine linkages in GAGs and can be directly labeled with fluorophores. WFA/VVA are primarily useful in fixed samples, not live tissue ^20^, and in living tissue may interfere with dynamics of learning-induced PNN modification. Additionally, CSPG expression and extracellular localization varies throughout the brain, and different CSPGs are differentially glycosylated, resulting in varying degrees of lectin binding and thus labeling ^14,21^. Imaging of ECM/PNN structures has been done using protein-specific antibodies, however antibodies are relatively large molecules that may not easily access epitopes in tight intercellular and perisynaptic spaces and can mainly be used in fixed samples.

Recent experiments using ChABC to disrupt PNN structures *in vivo* suggest that PNNs regulate experience-dependent synaptic plasticity ^11,12,22^ and stabilize drug addiction memories ^23^. Abnormal PNNs are correlated with deficits in memory; PNNs are disrupted in brain regions affected by Alzheimer’s disease and intact PNNs can protect neurons when they are exposed to toxic β-amyloid ^24,25^. Brain tissue samples from schizophrenia and seizure patients show PNN reduction in affected regions ^26,27^. PNN integrity and remodeling appears to be essential to normal brain function ^28^, similar to what is known about the broader ECM.

Based on the long-lived nature of some ECM/PNN molecules ^29,30^, we examined activity-dependent ECM/PNN changes in living tissues, hypothesizing that this stable and long-lived structures around neurons enables memories encoded in synapses to persist over time ^28,29^. To better characterize these structures, we first examine cultured neurons using both antibodies to proteoglycans and a HA-binding protein, used previously to study hyaluronan synthesis during neuronal development ^31^. We then attempted genetically encoded labeling; CSPGs are difficult targets for genetically encoded reporters due to their extensive posttranslational glycosaminoglycan modifications, however we tested fluorescent protein (FP) fusions to several other ECM proteins and identified the link protein *HAPLN1* (also called Crtl1) as a useful target for genetically labeling HAPLN1-containing extracellular structures both *in vitro* and *in vivo*. HAPLN1 is structurally and functionally important for ECM throughout the body, not just in the CNS. Our results describe ECM/PNN structure and establish a mouse model for imaging live ECM/PNN changes to better understand memory and the brain.

## Results

### WFA, aggrecan, and HA coat dendritic processes and surround synapses

ECM/PNNs are known to densely form around certain mature cultured cortical neurons (Fig 1A), so to first test basic structural labeling we used rat neurons grown in culture to image GAGs with WFA, HA with hyaluronic acid binding protein (HABP), and aggrecan with an anti-aggrecan antibody ^32^. Each label revealed a similar extracellular structure around neuronal soma and processes that had openings at PSD95-positive puncta, suggesting that these structures have the expected “holes” that were filled by mature synapses (Fig 1B-D). Cortical neurons in culture are a heterogeneous population, and only a fraction of neurons present possess extracellular aggrecan and WFA while the majority have some HA.

The well-known PNN label WFA is associated with structures that incorporate the heavily GAG-modified aggrecan ^33^, providing lots of binding sites for labeling, and we examined WFA labeling in adult mouse brain slices. The cortex showed a subset of heavily labeled neurons in cortical layers II-IV and VI, while in the cerebellum, WFA labeling appeared primarily in the granular layer of the folia. Several densely WFA-labeled neurons appear in the hippocampal CA2 region (Fig 2A). This observations are consistent with studies examining WFA labeling in mouse brain tissue ^34^. We found that aggrecan similarly surrounds a subset of neurons in cortical layers II-IV and VI, many Purkinje neurons in the cerebellum, and a larger subset of neurons in the hippocampus than those labeled with WFA (Fig 2B). There is some overlap between neurons positive for aggrecan and WFA in the cortex, and little detectable overlap between the cerebellum and hippocampus, considering the limitations of comparing different exogenously applied labels (Fig 2C). Together these data indicates that both WFA and aggrecan label individual neurons throughout the brain, and that these subsets overlap but are not identical.

**Figure 2.**
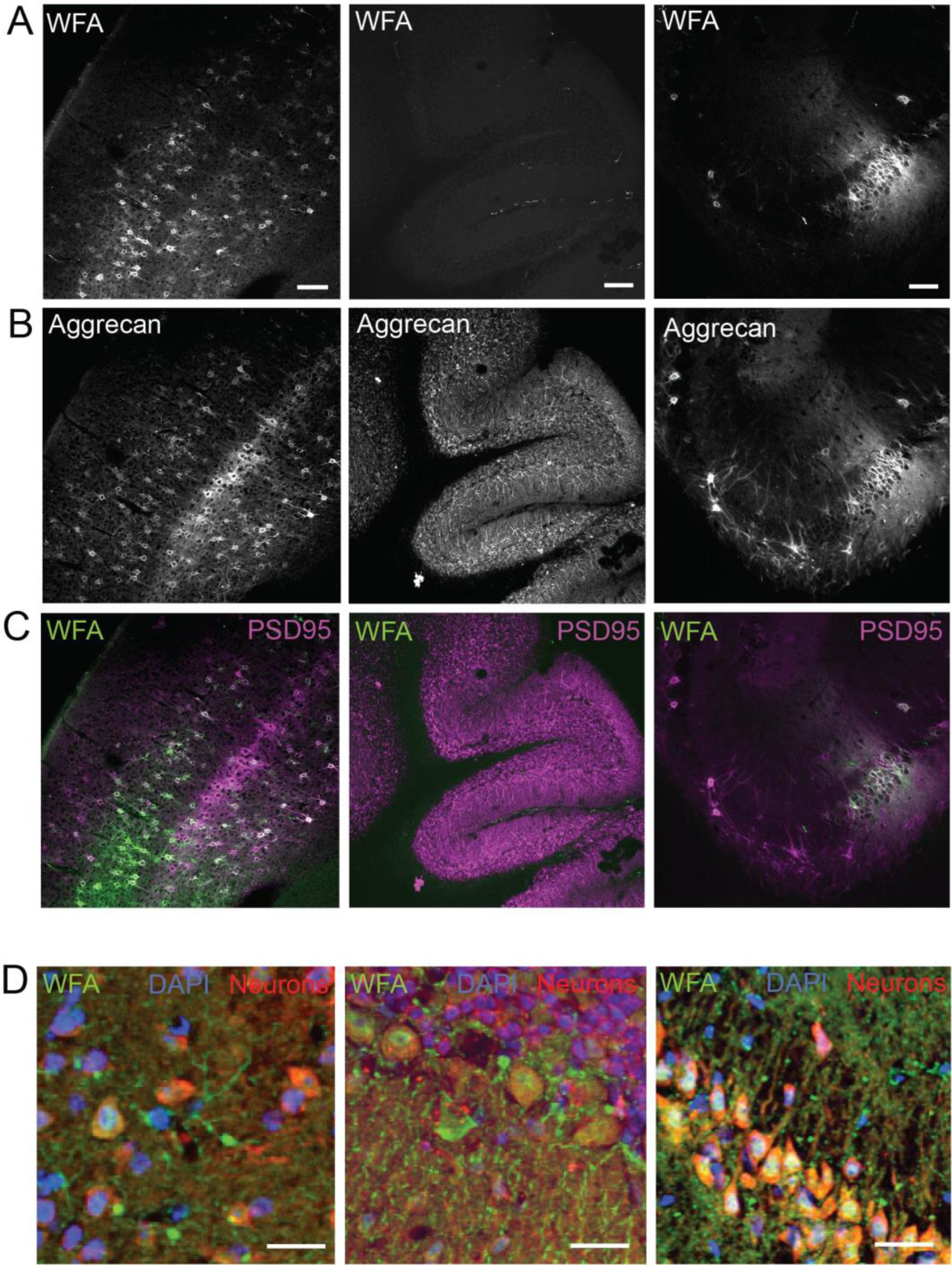
WFA and anti-aggrecan antibody label a small subset of neurons while most neurons are surrounded by HA. ***A***, Regions of mouse brain labeled with fluorescein-conjugated WFA. ***B***, Regions of mouse brain labeled with anti-aggrecan antibody. ***C***, Pseudocolor images with WFA labeling in green and anti-aggrecan antibody labeling in purple; WFA and aggrecan colocalize in some regions, but not all. ***D***, Pseudocolor images with HA labeling in green, nuclei in blue, and neurons in red; images show that all neurons in these regions are surrounded by a HA-based coating. Scale bars in ***A***-***C***, 20 μm; scale bars in ***D***, 20 μm.

Given that all PNNs are organized around an HA matrix, we expected HA to be a broader marker of both diffuse ECM/PNNs around neurons. It is known that the hippocampus contains less aggrecan and more versican and brevican^21^, which have fewer glycosyl modifications compared to other CSPGs and therefore may not be as strongly labeled with WFA. Using HA binding protein (HABP) in brain slices, we found that all brain regions express a dense layer of extracellular HA around neuronal soma and processes (Fig. 2*D*), suggesting that the extracellular structure visualized with WFA may appear in similar forms throughout the brain.

### HAPLN1-Venus is expressed, secreted, and integrated extracellularly around neurons

To investigate ECM/PNN involvement in plasticity and memory, we needed a method to monitor structural changes in living tissue. CSPGs are both difficult to label with FPs and differentially distributed in the brain, so we instead labeled the small HAPLN proteins that link CSPGs to HA via loop domains in their secondary structures ^35^. There are four known link proteins, HAPLN1-4, however HAPLN3 has little to no expression in brain tissue. HAPLN1 is the best characterized in animals; it is upregulated during critical period closure, present in all PNNs, and essential for proper PNN structure ^8,36^. Knocking out HAPLN1 in adult animals results in abnormal PNNs and memory deficits in adult mice, which makes it a functionally relevant target for neuronal ECM labeling ^36,37^. We fused HAPLN1, HAPLN2, and HAPLN4 to the FP Venus ^38^ and expressed fusions in primary cultured neurons (Fig 3A). HAPLN2 and HAPLN4 Venus fusion proteins were restricted to the cytosol, suggesting secretion was impaired following fusion to Venus. HAPLN1-Venus fusion protein however was produced, secreted, fluorescent, and integrated extracellularly around neurons.

**Figure 3.**
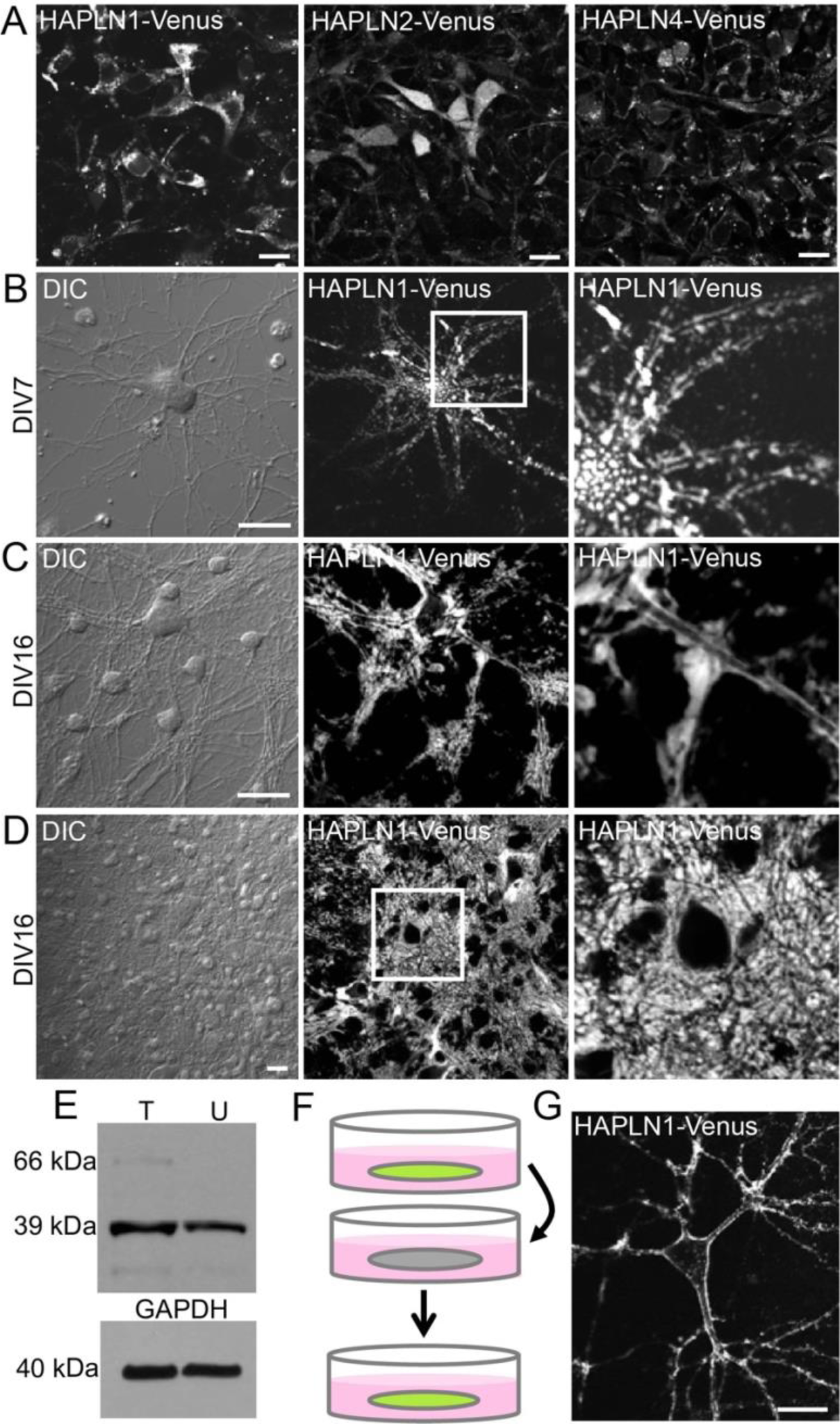
HAPLN1-Venus is expressed, secreted, and integrated extracellularly around neurons. ***A***, Cultured neurons transfected with a plasmid encoding each fusion construct; HAPLN1-Venus was expressed, secreted, and appeared to localize to the PNN. Nearly all HAPLN2-Venus and the entirety of HAPLN4-Venus remained intracellular. Scale bars, 20 μm. ***B***, Cultured neuron expressing HAPLN1-Venus at DIV7. ***C***, ***D***, Cultured neurons expressing HAPLN1-Venus at DIV16. ***E***, Western blot indicating that the level of HAPLN1-Venus fusion protein at 66 kDa is below the level of endogenous HAPLN1 at 39 kDa; T: transfected; U: untransfected. ***F***, Schematic describing experiment in which conditioned media from HAPLN1-Venus-expressing neurons was transferred to untransfected neurons. ***G***, Secreted HAPLN1-Venus fusion protein from conditioned media integrates extracellularly around untransfected neurons. Scale bars in ***A***, 20 μm; scale bars in ***B***-***D***, 20 μm; scale bars in ***G***, 20 μm.

We then monitored HAPLN1-Venus over time. By days in vitro 7 (DIV7), the fusion protein is expressed and bound extracellularly around neurons (Fig 3B) and localization is more uniform around neurons by DIV16 (Fig 3C, D). We confirmed that HAPLN1-Venus was not drastically overexpressed by Western blot (Fig 3E). Finally, we demonstrated HAPLN1-Venus secretion into the culture medium by transferring conditioned media from HAPLN1-Venus expressing neurons to untransfected neurons and observing extracellular HAPLN1-Venus integration around naïve neurons (Fig 3F, G).

To confirm that HAPLN1-Venus binds ECM in a HA-dependent manner, we treated neurons expressing HAPLN1-Venus with hyaluronidase, a bacterially derived enzyme that degrades HA, and monitored structural changes over time. HAPLN1-Venus fluorescence was diminished around neurons over a 10 h period after hyaluronidase treatment (Fig. 4A). We then co-labeled HAPLN1-Venus expressing neurons with a PSD95 antibody and found that HAPLN1-Venus surrounds PSD95 puncta, suggesting synaptic “holes” of similar structure as those found with WFA labeling and confirming that HAPLN1-Venus expression and integration permitted normal synapse formation (Fig 4B) and effectively allowed us to monitor dynamics of extracellular structures in culture.

**Figure 4.**
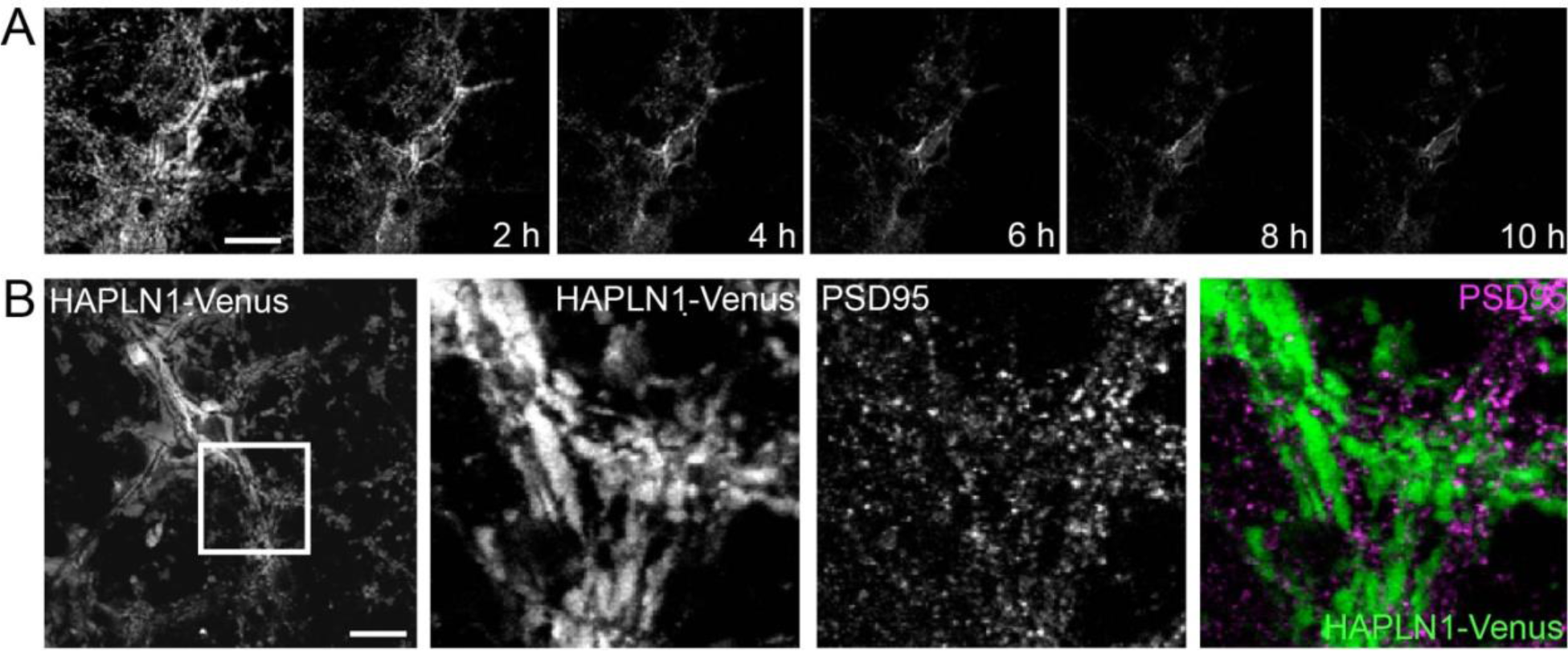
HAPLN1-Venus labels PNNs in living cultured neurons. ***A***, HAPLN1-Venus expressing neurons treated with hyaluronidase, which disrupted HAPLN1-Venus localization around neurons over 10 h. ***B***, HAPLN1-Venus is excluded from synapses labeled with an antibody to PSD95. Scale bars in ***A*** and ***B***, 20 μm.

We then asked whether HAPLN1-Venus fusion could label ECM *in vivo*. We generated a HAPLN1-Venus mouse knock-in (KI) via Crispr/Cas9 genome editing (Janelia) (Fig 5A-G). These KI animals show Venus fluorescence in all tissues that endogenously contain cartilage and connective tissue (Fig 5A-C). HAPLN1-Venus is clearly expressed in the brain (Fig 5D, E) and exhibits a dense, fenestrated structure characteristic of PNNs around many neurons (Fig 5F, G). This HAPLN1-Venus KI model can be used to further study ECM and PNN *in vivo* and may help us better understand the functions of extracellular changes at synapses over time.

**Figure 5.**
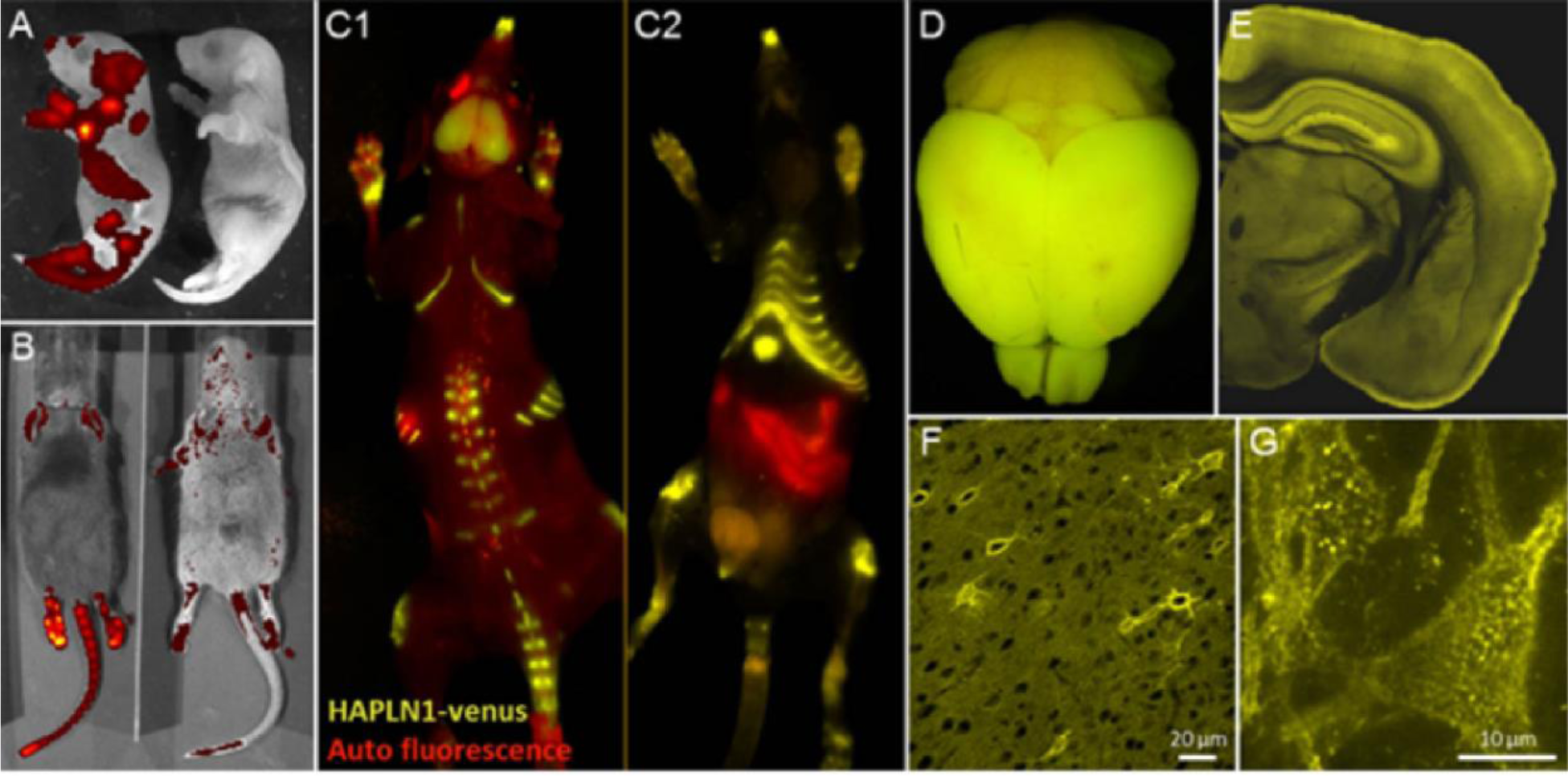
HAPLN1-Venus is expressed in a living knock-in mouse, localizing to brain and cartilage. **A**, HAPLN1-Venus transgenic mouse newborn pup on the left and a wild-type C57BL/6J mouse newborn pup on the right. **B**, HAPLN1-Venus fluorescence in a homozygous (right) and heterozygous (left) knock-in animal. **C1, C2**, Two views of HAPLN1-Venus fluorescence in cartilage. **D, E**. HAPLN1-Venus fluorescence in mouse brain. **F, G**. HAPLN1-Venus fusion protein localizes more heavily around some neurons than others and shows punctuated localization around neurons that express more HAPLN1-Venus. Scale bars in **F**, 20 μm; scale bar in **G**, 10 μm.

## Discussion

For as long as PNNs were thought to be primarily around PV+ inhibitory interneurons in the cortex ^39^, it was hard to imagine a wider role in learning and memory. Here we imaged traditional PNN structures using lectin labeling, but additionally showed that an HA-based, PNN-like ECM is more widespread and present around many neurons in the CNS, consistent with how information is stored in areas throughout the brain. We describe the HAPLN1 link protein as a target for genetically encoded labeling, elucidating structures around both traditional PNNs and the broader ECM, and describe an animal model for monitoring the dynamics of these extracellular structures in living animals using microscopy.

Genetically encoding reporters into the ECM/PNN via HAPLN is a helpful strategy because it enables us to track these structures in living brain tissue. Previously, lectin labeling *in vivo* had little success and was restricted to imaging specific points in time ^20^. Genetically encoded labeling was a logical next step, however CSPGs are post-translationally modified, making them difficult targets for fusions ^40^. HAPLN1 fusions are a more straightforward way to target ECM/PNN in live tissues, based on how HAPLN1-Venus can be produced, secreted, and integrate extracellularly in cultured neurons and *in vivo*. Further studies using our HAPLN1-Venus knock-in animal for monitoring structural changes in combination with genetically encoded reporters such as miniSOG or APEX2 for correlated light and electron microscopy ^41–43^ will be invaluable in understanding how the ECM/PNN is involved in memory encoding and consolidation. Considering how HAPLN1 is expressed in ECM throughout the body and also appears within cartilage, our mouse model may prove to be a useful tool for a broader field of studies on ECM structure and integrity throughout the body.

HAPLN1 is emerging as an important extracellular molecule implicated in memory, independent of our use as a target for genetically encoded labeling in the ECM. Disrupting PNNs in a HAPLN1 (Crtl1) mouse knock-out model rendered fear memories susceptible to erasure ^44^. HAPLN1 turnover was found to be relatively rapid in PNNs and at synapses ^45^, and viral-mediated HAPLN1 delivery into mouse cortex showed that HAPLN1 is both necessary for PNN maturation in mice and when overexpressed can accelerate PNN and memory engram formation ^46^. Further studies on HAPLN1 modulation in the brain will help understand how it may function in memory encoding and persistence.

We hypothesized that synaptic changes made to store a long-term memory should be independent of the need to copy information repeatedly ^28^. Many intracellular proteins face relatively rapid turnover, being synthesized and degraded on the scale of hours to several days ^47,48^. These are susceptible to cellular or brain-wide disruptions that do not necessarily affect memory, including seizure, barbiturate overdose, or metabolic stress, and we speculate that they may be unreliable substrates for very long-term information storage. Several synaptic proteins have recently been implicated in memory maintenance, including PKMζ, CPEB, CaMKII, and Arc/Arg2.1, but these proteins have only been examined in relatively short-term memory models and turn over on the time scale of hours to days ^49–52^. ECM molecules can be much more stable than intracellular proteins and remain sequestered away from intracellular ubiquitin and enzyme-mediated degradative processes, and thus are thought to be less susceptible to disruption. ^30,47^

Our results allow us to now ask new questions about ECM/PNN in the brain. How stable are extracellular structural changes produced when a synapse is strengthened? How is the PNN modified at synapses modulated by learning and memory extinction? Are they modified when a memory is recalled? We have yet to understand the significance of variations in ECM/PNN density and composition, which may be understood better using genetically encoded labeling and imaging in the extracellular space. It is now possible to experimentally address these questions in living animals using our HAPLN1-Venus mouse model.

Here we report a new tool for monitoring dynamics of the ECM/PNN, extracellular scaffolds that may be differentially composed but nevertheless exist in some form around all neurons. Further studies combining labeling and imaging methods to visualize dynamics in living animals will help us better understand the role of these elusive structures in the brain.

## Materials and Methods

### Reagents

Mouse HAPLN1 cDNA was purchased (Origene, Rockville, MD) and cloned as a fusion to Venus into the pCAGGS mammalian expression plasmid, which drives expression via a cytomegalovirus (CMV) enhancer fused to a chicken beta-actin promoter. This construct was generated using standard molecular biology techniques; PCR, restriction enzyme digestion, and ligation. All subcloned fragments were fully sequenced to confirm successful construction. Primary antibodies used were mouse monoclonal anti-PSD95 (Pierce, Rockford, IL), rabbit polyclonal anti-Aggrecan (Millipore, Darmstadt, Germany), rabbit polyclonal anti-Synapsin I (Millipore), mouse monoclonal anti-GFAP (Sigma-Aldrich, St. Louis, MO), rabbit polyclonal anti-HAPLN1 (GeneTex, Irvine, CA). For immunoblotting, primary antibodies were used at 0.1-0.4 μg/mL. HRP-conjugated mouse secondary antibody (Cell Signaling, Danvers, MA) and HRP-conjugated rabbit secondary antibody (BioRad, Irvine, CA) were used at 0.4 μg/mL. For immunofluorescence, primary antibodies were used at 0.5-1 μg/mL. Alexa Fluor 568-conjugated mouse secondary antibody and Alexa Fluor 647-conjugated goat secondary antibody (Life Technologies, Carlsbad, CA) were used at 0.5 μg/mL. Streptavidin Alexa Fluor 488, Streptavidin Alexa Fluor 647, and unlabeled Streptavidin (Life Technologies) were used at 2 μg/mL. Fluorescein-conjugated Wisteria floribunda agglutinin and Vicia villosa agglutinin (Vector Labs, Burlingame, CA) were used at 10 μg/mL. Biotinylated hyaluronic acid binding protein (Millipore) was used at 10 μg/mL. For time lapse imaging, hyaluronidase (Sigma) was used at

### Cell culture

All cell culture reagents were obtained from Life Technologies unless otherwise indicated. Cortical neurons were dissociated by papain from postnatal day 2 (P2) Sprague Dawley rats of either sex, transfected by Amaxa electroporation (Lonza, Basel, Switzerland), then plated on poly(D-lysine)-coated MatTek glass-bottom dishes (MatTek Corporation, Ashland, MA) and cultured in Neurobasal A medium with B27 supplement, 2 mM GlutaMAX, 20 U/mL penicillin, and 50 μg/mL streptomycin as previously described ^53^. All neuron experiments described were performed in cultured neurons matured to between 14 and 18 days in vitro (DIV) unless otherwise specified. All animal procedures were approved by the Institutional Animal Care and Use Committee of the University of California, San Diego.

### Immunoblotting

For analysis of the expression level of HAPLN1-Venus, transfected neurons were plated and maintained in six-well cell culture plates. At DIV14, neurons were rinsed quickly with ACSF (in mM): 125 NaCl, 2.5 KCl, 1 MgCl_2_, 2 CaCl_2_, 22 D-glucose, and 25 HEPES (pH 7.3) and resuspended in lysis buffer (20 mM Tris-HCl, 1% Triton detergent, 10% glycerol, 2 mM EDTA, 137 mM NaCl, and Roche EDTA-free protease inhibitor, pH 7.4). Cells were homogenized and protein content assessed by Bradford assay. SDS loading buffer was added to lysate, then immunoblotting was performed. Briefly, lysates were run on NuPage 4-12% Novex Bis-Tris SDS polyacrylamide gels (Life Technologies). Proteins were transferred onto polyvinylidene fluoride membranes (Life Technologies) by electroblotting, and membranes were then blocked with 10% non-fat dried milk in Tris-buffered saline with 0.1% Tween-20 (TBST), incubated in primary overnight in 5% BSA in TBST at 4°C, washed in TBST to remove excess primary antibody, incubated in HRP-conjugated secondary antibody in TBST for 1 h at room temperature, and finally rinsed 3 times for 15 min each in TBST. Proteins were visualized by chemiluminescence using SuperSignal West Pico Chemiluminescent Substrate (Thermo Scientific) and film (Blue Devil, Genesee Scientific).

### Time-lapse microscopy

Neurons were imaged on an Olympus FluoView 1000 Confocal LSM equipped with a stage-top environmental chamber with temperature set to 37°C and 5% CO_2_, using a 40x oil objective. For confocal imaging, the following settings were used: excitation with a 488-nm argon-ion laser line at 0.5% power, scan resolution 800 x 800 pixels, scanning speed 2 μsec/pixel, photo-multiplier voltage 700 V, digital gain of 1. A stack of optical sections at 1-μm intervals through each neuron was obtained at each position and time point and then flattened in a maximum intensity projection. Analysis was performed on an Apple Macintosh notebook using the US National Institutes of Health ImageJ program. Immunofluorescence and time lapse images of related conditions were matched for laser power, gain, and offset.

### Immunofluorescence in cultured cells

Neurons were fixed by addition of one culture volume of 4% paraformaldehyde and 4% sucrose for 10 minutes at room temperature (RT), then washed with phosphate-buffered saline (PBS), blocked in PBS with 5% non-immune goat serum (Sigma), and probed for PSD95 following standard procedures. Specificity of secondary antibodies was confirmed in control samples without primary antibody. Images were obtained on an Olympus FluoView 1000 Confocal Laser Scanning Microscope (LSM) using a 40x oil immersion lens. A stack of optical sections with 1024 x 1024 resolution at 1-μm intervals through each neuron was obtained and then flattened into a maximum intensity projection. Analysis was performed on an Apple Macintosh notebook using the National Institutes of Health ImageJ Program. Immunofluorescence images of related conditions were matched for laser power, gain, and offset.

### Immunohistochemistry

Male FVB mice aged 6 months were anesthetized by intraperitoneal administration of Ketamine (100 mg/kg; Ketaset, Pfizer, New York, NY) supplemented with Midazolam (100 mg/kg; Midazolam, Akorn, Lake Forest, IL) and transcardially perfused first with 50 mL of oxygenized Ringer’s solution and then 100 mL of 4% PFA in DPBS. The brain was dissected and post-fixed in 4% PFA for 1 h at 4^0^C, then cryoprotected in DBPS buffer containing 30% sucrose overnight. Brain tissue was embedded in O.C.T. (TissueTek, Radnor, PA) then cut on a cryostat into 10 μm-thick sagittal sections and mounted onto glass slides. Primary antibodies were incubated overnight at RT in PBS containing 0.1% BSA, 0.1% fish gelatin (Sigma, Cat #), and 0.1% Triton X-100. Sections were then incubated with secondary antibody for 2 h at RT. After washing with PBS, samples were mounted in VectaShield Mounting Medium with DAPI (VectorLabs, Burlingame, CA) and imaged on an Olympus FluoView 1000 Confocal Laser Scanning Microscope (LSM) using a 40x oil immersion lens. Images were taken at 1024 x 1024 resolution.

### Transgenic mouse generation

The HAPLN1-Venus transgenic mouse was generated by Crispr/Cas9-mediated knock-in of mouse HAPLN1 fused to Venus via the C-terminus, conducted by the Gene Targeting and Transgenics Facility at the Janelia Research Campus in Ashburn, VA. HAPLN1-Venus transgenic mice are available via The Repository at the Jackson Laboratory (JAX Repository), Stock No. 032115.

## Author Contributions

SPL and VLR performed experiments; SPL, VLR, MHE, and RYT contributed to the design of the experiments and analyzed data; SPL, VLR, RYT, and MHE wrote the manuscript.

## Acknowledgments

We would like to thank John T. Ngo, John Y. Lin, and Stephen R. Adams for useful discussions and suggestions, and Daniela Boassa, Mason R. Mackey, Hiro Hakozaki, Eric A. Bushong, Andrea Thor, and Thomas Deerinck for assistance with microscopy and analysis. This work was supported by NIH grants awarded to RYT, VLR, and MHE (NS027177), to RYT and MHE (R01 GM086197), and to MHE (P41 GM103412). SPL was supported by funding from the USCD Graduate Training Program in Cellular and Molecular Pharmacology (T32 GM007752) and the UCSD Graduate Training Program in Neuroplasticity of Aging (T32 AG000216).

## Notes

### Competing Interest Statement

The authors have declared no competing interest.

